# Sub-cellular dynamic investigation of the multi-component drug on the gastric cancer cell BGC823 using Raman spectroscopy

**DOI:** 10.1101/2022.03.24.485691

**Authors:** Wenhao Shang, Teng Fang, Anpei Ye

## Abstract

The potential of Raman spectroscopy in anticancer drug study has been demonstrated, yet its ability to character systematic cellular changes caused by multi-component drugs has not been explored. Here we used micro-Raman spectroscopy combined with bright field imaging to study Compound Kushen injection (CKI) at a sub-cellular level including intracellular vesicles(IVs). In our report, CKI caused dysfunction of DNA replication and repair was displayed by Raman spectrum (RS) from the cell nucleus. Meanwhile, the dynamics of CKI induced intracellular vesicles and cell component deconstruction was delineated by RS from the cytoplasm and IVs. The lipids-related biomolecular changes were also presented by the cytoplasm RS: the lipids level in the cytoplasm first descended then uprising. In conclusion, this study validated the mechanism and displayed the dynamics of CKI in treating cancer cells. We proved the capability of subcellular micro-Raman spectroscopy for detecting systematic cellular changes and its application for multi-component drug evaluation.

---

“Multiple component-therapeutics” anticancer drugs such as Traditional Chinese Medicines (TCMs) have gained more and more attention in drug discovery^1^. The main advantages of TCMs are their function on improving the efficacy of cancer therapy and reducing side effects and complications^2, 3^, also on modulating immune function and improving the quality of life of cancer patients in clinical use^4^. Compound Kushen injection (CKI) is a National Medical Products Administration approved TCM formula used in the clinical treatment of various types of cancers in China^5^The chemical fingerprint of CKI contains at least 8 different components, with primary compounds Matrine and Oxymatrine^6^. It has been shown multiple bioactive ingredients in CKI deliver an integrated anti-tumor effect through multiple targets and their associated molecular pathways^7^. CKI is proved to suppress cell cycle and DNA repair pathways, even reducing the metabolism level in cancer cells^8^. Studies proved that Matrine could inhibit cell proliferation and introduce apoptosis in various cancer types via different molecular pathways^8, 9^. In human HepG2 cells, Matrine induced autophagy in a dose-dependent manner^10^ In short, the existence of CKI multiple bioactive ingredients causes multi-level cellular changes from morphology to DNA replication/repair inhibition, cell proliferation inhibition, autophagy, and apoptosis^1, 8, 11, 12^.

Micro-Raman spectroscopy (RS), known as molecular fingerprint spectroscopy, is a label-free and noninvasive technique to characterize the chemicals component and content in cell samples^13, 14^. The application of RS in drug screening and investigation of cell response profiles has been explored in many anticancer drugs such as cisplatin (an alkylating and DNA binding agent), doxorubicin, vincristine, paclitaxel ect^15-18^. Some of the studies even explored the study of the anticancer drug at the subcellular level^15, 19-21^. However, the potential of RS for multi-component drug study has not been explored for its complex cell response. Meanwhile, the drug-induced intracellular vesicles (0.4-1um) activity has not been investigated at sub-cellular RS study, due to the difficulty of acquiring RS with a high resolution.

In this report, we used CKI as a demo for demonstrating RS potential for multi-component drug study at the subcellular level including intracellular vesicle activity. Using a custom-built 532nm laser Raman platform with a high-NA (numeric aperture) objective (100×/ 1.46) enables us to characterize the dynamics of cell intracellular vesicles-related cell activities at around 200nm resolution. We used CKI and 5-fluorouracil (5Fu, as a reference) to treat gastric cancer cell line BGC-823 at different drug concentrations and time points. First, Cytotoxicity assays and Trypan blue cell counting was conducted to verify CKI effects on cell proliferation and viability inhibition, and also identified the equivalent cytotoxicity effect concentration of CKI and 5FU. Subsequently, the nucleus RS of cells was collected after CKI 5Fu was treated for verifying the CKI induced DNA replication/repair and proliferation inhibition. To delineate the dynamics of CKI induced intracellular vesicles, cell images, cell cytoplasm, and vesicles RS signals of the same cell were collected at the same time.

## EXPERIMENTAL SECTION

### Cell culture and drugs

CKI with a total alkaloid concentration of 20.4 mg/ml in 5 ml ampoules, human BGC823 gastric carcinoma cells, and 5-Fluorouracil were provided by Beijing Cancer Hospital. The cell culture method has been described in our previous work^14^. In short, BGC-823 cell was placed in standard culture medium: RPMI-1640 medium (Macgene, Hangzhou, China) supplemented with 10% fetal bovine serum (Tianhang Biological Technology Co. Ltd., Beijing, China) with antibiotics and cultured at 37 °C with a relative humidity of 95% and 5% CO_2_. After cells reached 40% confluence the cell culture was replaced by drug mixed ones.

### Cell viability and Live-cell counting

The cholecystokinin (CCK-8) assay experiment was conducted to evaluate the anticancer effect of CKI. The wells of 96-well trays were seeded with 1×10^4^ cells suspended BGC-823 cells in 100 μL of medium and cultured overnight. Next, we cultured and tested the cell viability treated by CKI(2mg/ml,1mg/ml) and 5-FU (10ug/ml) for 24, 48, and 72 hours, following the procedure described previously^14^. In the meanwhile, live-cell counting was performed in parallel with RS data collection using a hemocytometer since the dead cells were marked by trypan blue but the living cells were not^22^.

### Sub-cellular Raman spectroscopy & bright filed cell imaging

We constructed an optical configuration for the 532nm laser stimulated back-scattering RS collection and cell imaging as described previously^23, 24.^ Especially, a quite high-NA (numeric aperture) objective (100×/ 1.46 oil, N-Achroplan, Zeiss, Oberkochen, Germany) was used herein to achieve sub-cellular high-resolution Raman Spectroscopy. The size of the laser spot on the cells can be estimated by the Bassel function for a Gaussian laser beam, Dmin=1.22λ/NA, where λ=532nm is the wavelength of the laser, and Dmin is the diameter of the Airy disc which contains 84% of the whole laser beam energy^25.^ In our experiment, the theoretical value of Dmin was about 444nm. Normally, the size of BGC823 cells ranges from 10μm to 20μm, and the size of observed intracellular vesicles (IVs) are from 0.4um to 0.9um. Thus, we were able to detect RS signals from the sub-cellular structures. It should be noted that with the high NA objective the system could only achieve high spatial resolution in the x-y plane, not in the z-direction. Therefore, the RS signal of vesicles inevitably includes the contribution from the cytoplasm. However, since RS signal intensity is directly proportional to the excited power, and the light strength is sharply decreased outside the focal point of the laser when the laser was precisely focused on vesicles, most of the RS signal we collected come from the IVs rather than the cytoplasm around the vesicles. Thus, we could approximately see it as the RS of IVs. In our experiment, the laser power was 14 mW at the focal plane of the objective and the integration time was the 30s for each spectrum acquisition. Meantime, we collected the bright-field images for each cell synced with RS measurements. As a result, more than 25 cells were collected for each different condition (Untreated, CKI 1mg/ml, CKI 2mg/ml, 5Fu 10ug/ml), respectively.

The RS analysis was performed using OriginPro (2019b, OriginLab Corporaton.US) and in-house scripts based on the R (3.6.1), and MATLAB (2019b, The MathWorks, Inc. US). For each spectrum, the cosmic rays were firstly removed from raw data, then 3rd-order Polynomial Curve Fitting was conducted to remove the background envelopes followed by smoothing with 5-point Savitzky-Golay, the spectral region from 500 cm^-1^ to 1800 cm^-1^ that contained abundant biomedical signals remained. All the RS was normalized by area. As our previous work showed^16^, the area under the RS curve is a more suitable and accurate index.

## RESULTS & DISCUSSIONS

### Inhibition of cellular proliferation and viability

MTT assays (CCK8) were conducted to measure cell viability after treating with different doses of CKI at 24h, 48h, 72h to quantitatively validate the effect of CKI on cell proliferation in gastric cancer BGC-823 cell line. Figure 1. a attests that the cell viability of BGC-823 cells was significantly inhibited by a high dose of CKI (2mg/ml, based on the total alkaloid concentration in CKI) and 5Fu(10ug/ml). The cell numbers were counted by Trypan blue staining (only live cells were counted). Figure 1. b shows that the live cell numbers were greatly decreased by CKI 2mg/ml and 5Fu 10ug/ml. As shown above, CKI worked in a dose-dependent manner which is consistent with the previous reports^26.^ In short, the results validated CKI inhibited proliferation and viability of BGC-823 cells, and displayed CKI 2mg/ml and 5Fu 10ug had an equivalent cytotoxic effect. It should be noted that the concentrations of CKI are very high (in mg/ml range) and out of the physiological range. Here we just highlight the effects of the drug.

**Figure 1:**
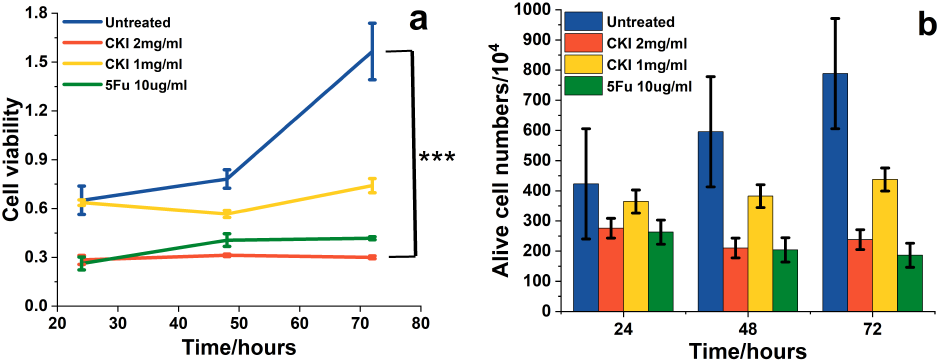
CKI inhibited proliferation and cell viability. a) Inhibition of BGC823 cell viability with CKI. The viability was measured by CCK8 kit (MTT). b) The numbers of different drug-treated conditions were counted after Trypan Blue stain. Data are represented as mean ±SEM. a, Two-way ANOVA ***<0.01.

### The nucleic acid decrease in the cell nucleus

The cell nucleus area, containing most of the cell DNA and in charge of RNA (mRNA, tRNA, rRNA) synthesis, is the main target of anticancer drugs. 5Fu exerts its anticancer effects through inhibition of thymidylate synthase and incorporation of its metabolites into RNA and DNA then disrupting normal DNA and RNA processing and function and the inhibited thymidylate is necessary for DNA replication and repair^27^. CKI can increase the level of DNA double-strand breaks (DSBs) and inhibit DNA repair and replication^28-30^. We used the RS from the cell nucleus to validate the drug effect on nucleic acid components. By comparing the RS alterations(figure 2), the difference of cells after CKI and 5Fu treatment were displayed by those RS peaks related to nucleic acids and other cell components. In eukaryotic cells, the 794 cm-1, 941 cm-1,1092 cm-1 and 1579 cm-1band of RS corresponds to nucleic acid components^31^. The characteristic peaks are assigned in Table (Table S1) based on previous studies involving several cell lines and biomolecules.^31-36^ Figure 2 and Figure S3 shows that the Raman spectra profile of treated groups were decreased after 24h, which mainly reflected in those Raman bands assigned to nucleic acid: 794 cm^-1^(DNA), 941cm^-1^(RNA), 1092 cm^-1^(DNA), 1375 cm^-1^(A, G, T) 1579 cm^-1^(DNA). The changes at 48h shown in figure S1 were similar to those at 24h. The decrease of DNA and RNA components at the nucleus validated CKI and 5Fu effect on cell DNA and RNA.

**Figure 2:**
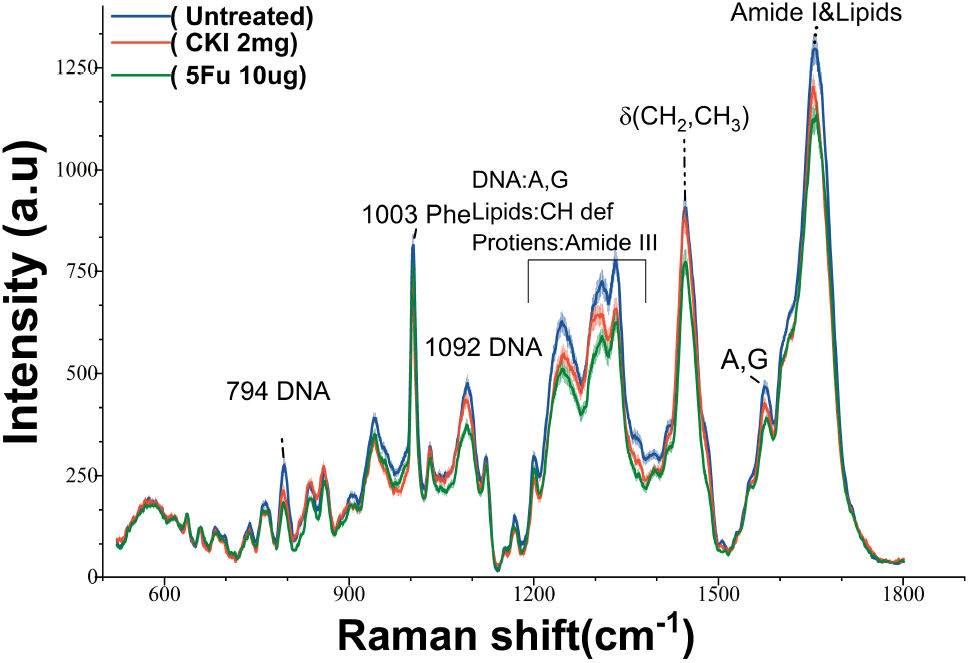
Nucleus area RS in BGC-823 treated with CKI or 5F. the Nucleus RS at 24h (mean ±s.e.m., the light shadow represents the s.e.m)

To quantitatively compare the difference of nucleic acid reduction between CKI and 5Fu, we compared the 794 cm^-1^ intensity and the area under the peaks from 1287-1343 cm^-1^ (mainly reflected the A C G in nucleic acid). Pair-wise comparisons involving more than two groups were evaluated using the appropriate Bonferroni corrections and a one-way ANOVA test was conducted with p<0.05. Both CKI and 5Fu had significantly reduced those nucleic acid signals. While 5Fu had a stronger nucleic acid inhibition effect, which is understandable since 5Fu inhibits thymidylate synthase and incorporates its metabolites into RNA and DNA rather than only involving in DSBs and DNA repair.

Additionally, the strong signal at 1653 cm-1, encompassing contributions from protein v(C=O) (amide I), lipids v(C=C), decreased in intensity after the drug-treated (Figure. 2 e). The proteins and lipids decrease also confirmed by characteristic peaks at 1003 cm-1(Phenylalanine), 1313 cm-1(carbohydrates) 1334 cm-1(CH3/CH2 wagging, protein), 1446 cm-1(CH2 bending). Surprisingly, for CKI-treated groups at 48h the relative intensity increased, and its position at 1656 cm-1,1454 cm-1, 1304 cm-1,1092 cm-1 was slightly shifted(Figure S1), which should be resulted from the increase of nearby lipids bands.^35^These RS changes in DNA and RNA components from the cell nucleus validate CKI and 5Fu effect on the nucleic acid.

### Intracellular vesicles accumulation

Many studies show that CKI can induce cell apoptosis^5, 8, 29^, which is characterized by a series of common morphological and biochemical features that include cell shrinkage, membrane blebbing, nuclear condensation, DNA fragmentation, mitochondrial fragmentation^37^. CKI was also shown to cause cell autophage^10^, characterized by the appearance of a double-or multi-membrane cytosolic vesicle for degradation of the cell component^37-39^. Moreover, autophagy-triggered cell death as autophagy is, strictly speaking, a mode of cell survival, but persistent autophagy generally triggers apoptosis^40^

Thus, the monitor of the intracellular vesicles (IVs) and related cytoplasm dynamics would be one of the keys to investigating the CKI anticancer effect. Due to the small size (0.4-1um) of IVs, a high-NA objective(100X/1.46) was applied to acquire high-spatial-resolution RS(around 200nm, see method for details). For detecting the cell morphology changes and IVs activity, we collected plenty of cellular photographs under the bright-filed imaging and measured corresponding the RSs of the IVs and cytoplasm for the same cell in different treatment conditions and time points, respectively.

The remarkable morphological changes were observed under CKI treated groups both at 24h and 48h (figure 4 b, e; figure 5 b). Abundant cytoplasmic vesicles (pointed out by white arrows in figure 5 b) with varying sizes (0.4-1um) were only observed in BGC823 cells treated by CKI. However, the cell size did not remarkably change. As for 5Fu-treated BGC823 cells, the size was larger than those of the untreated group(figure 4 c,f), which consists of our former work^41^.

**Figure 3:**
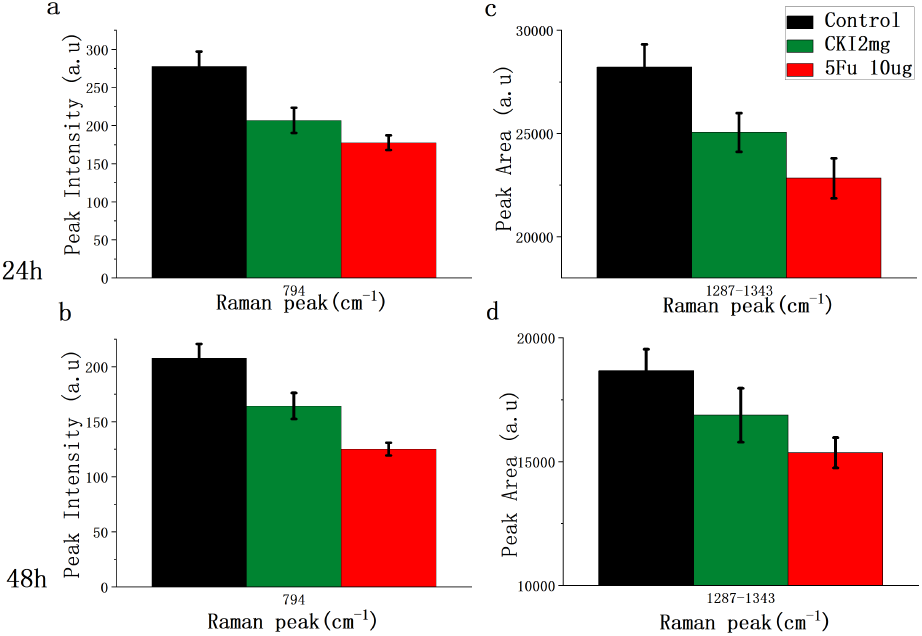
Bar plot of RS at 794 cm^-1^ and 1297-1343cm^-1^ a,b) The RS intensity at 794 cm^-1^ of different groups(control black,CKI 2mg green,5Fu10ug red)at 24h and 48h; c,d) The area under spectrum from 1297cm^-1^ to 1343cm^-1^different groups at 24h and 48h. Mean ± s.e.m., (the three groups were compared by a one-way ANOVA, and *P* < 0.05 in each figure)

**Figure 4.**
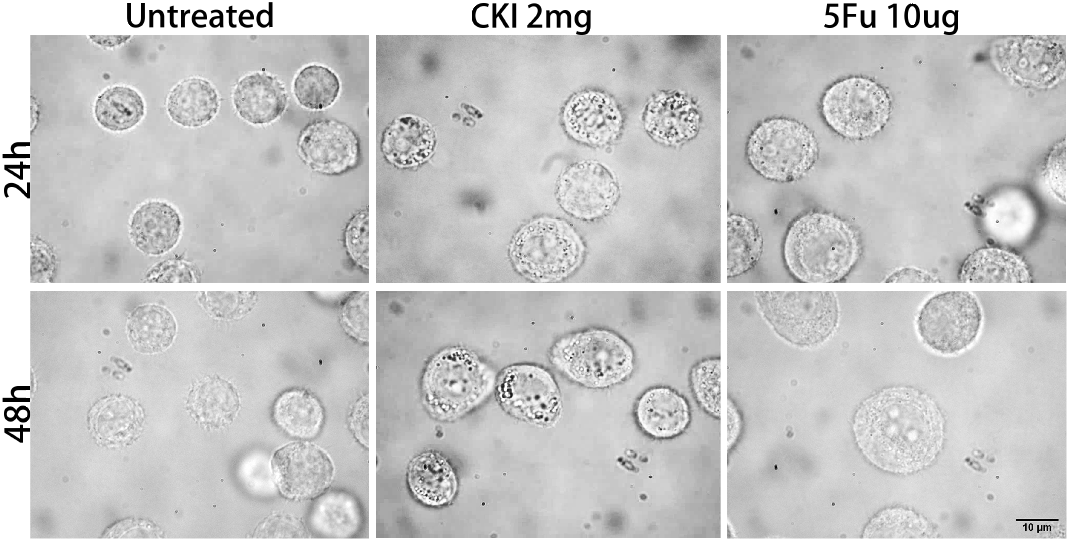
Cell morphology. a-c) The images of BGC823 cells after CKI/5Fu treatment for 24h in bright field; e-f) The images of BGC823 cells after CKI/5Fu treatment for 48h.

**Figure 5.**
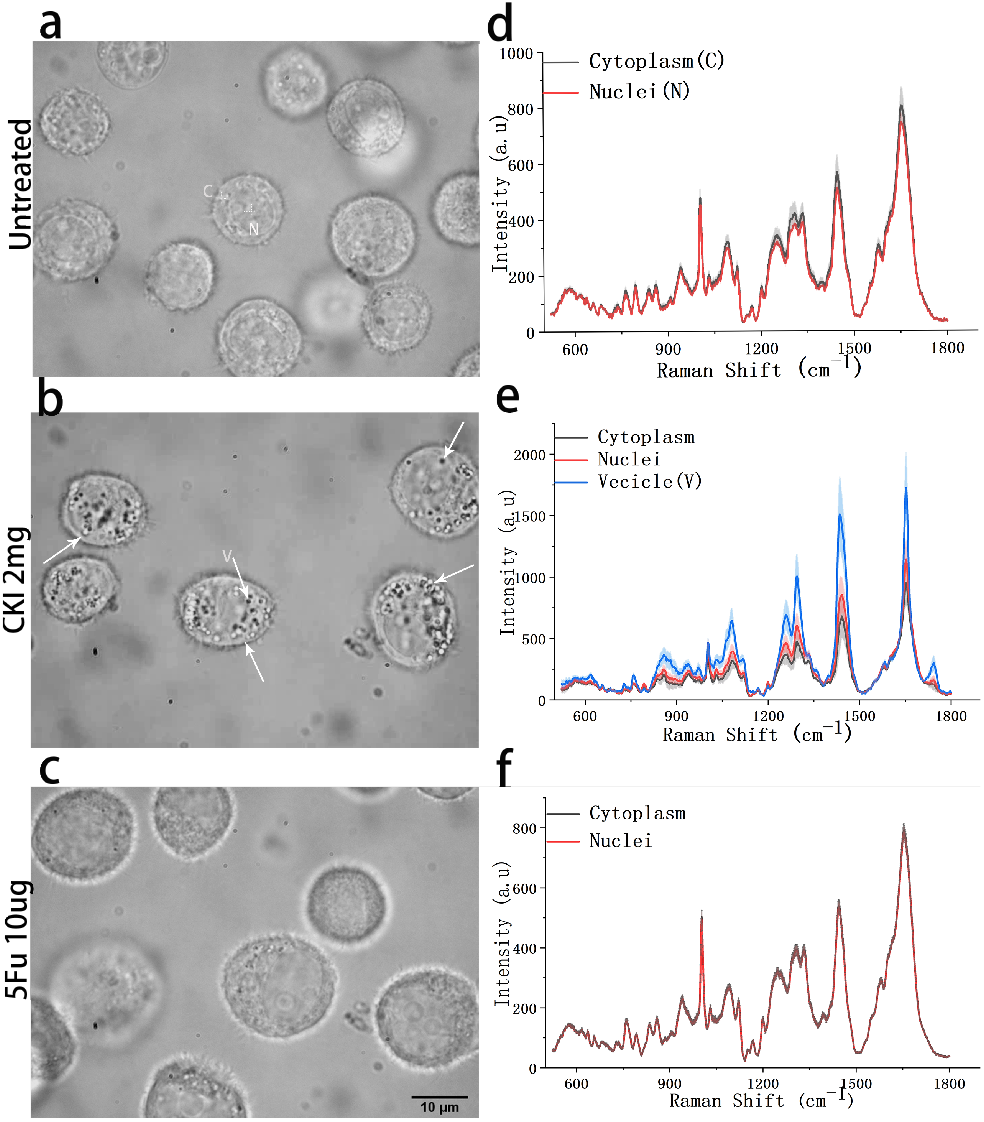
Cell images and subcellular RS of BGC-823 cells. a, d) The image and sub-cellular RS of the same cell without drug-treatment; b, e) The image and sub-cellular RS of the same cell under CKI 2mg/ml treatment; f) The image and sub-cellular RS of the same cell under 5Fu 10ug/ml. Note: all these data were collected after treatment for 48 hours, all RS were presented by mean±s.e.m., the light shadow represents the s.e.m

**Figure 6.**
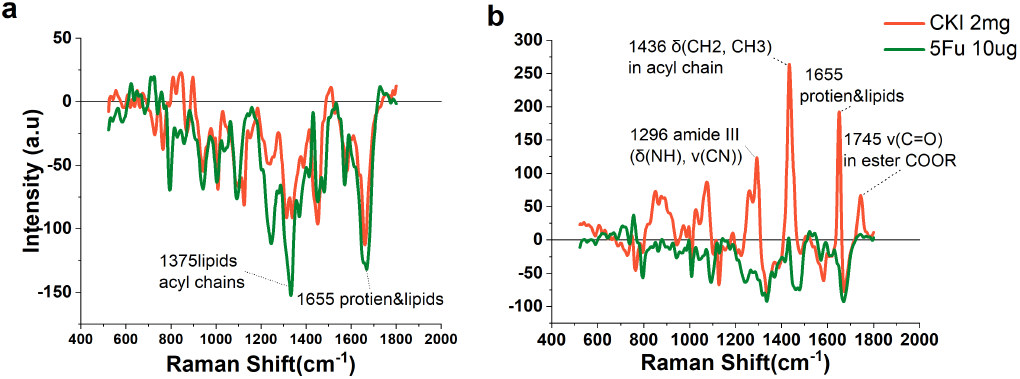
Cytoplasm RS of BGC-823 cells treated with CKI or 5Fu. a) the RS of drug-treated cytoplasm RS subtracting the untreated cytoplasm RS at 24h; b) the RS of drug-treated cytoplasm subtracting the untreated one at 48h.

### RS of Intracellular vesicles and cytoplasm

For analyzing the composition of enormous IVs and biochemical alteration in the cytoplasm, the RS from intracellular vesicles, cytoplasm, and nucleus of the same cells were analyzed. We compared the difference of RS between the Cytoplasm and nucleus, only in the CKI treated group the RS signals from the cytoplasm were significantly different and lower than the nucleus (figure 5 d-f). The RS from IVs had strong signals exactly at the peaks that the cytoplasm decreased (figure 5 e). The results from at 24h were the same as at 48h(figure S4). This indicated that the components fluxing from the cytoplasm to vesicles occurred within the CKI-treated cells. This was consistent with the cell component degradation process undergoing during apoptosis and autophagy.

The main peaks of vesicles appear in 1746 cm^-1^ (COOR), 1656 cm^-1^ (proteins), 1439 cm^-1^(carbohydrates/lipids), 1296 cm^-1(^lipids), 1199 cm^-1^ (proteins), 1081 cm^-1^(proteins/lipids/glycogen), 1030 cm-1(proteins/lipids/glycogen). Wherein the new peaks of 1746 cm^-1^ reflected that a mass of phospholipids was used to construct vesicles. The protein, RNA, carbohydrates, and lipids signals were enhanced due to deconstructed cell components during autophagy and apoptosis. We also collected the sub-cellular spectra for the CKI-treated 72h-group, almost all of the spectra features (Figure S2) are the same as those of the 48h-group. The only difference was at 1746 cm-1(COOR), 72h-group did not strong increase like 48h-group. This band is the key difference between lipids and fatty acids in cell components. It suggested fatty acids rather than lipids kept increasing in the cytoplasm after CKI treatment.

To further analyze the cytoplasm alterations companied with IVs accumulation, we used drug-treated cytoplasm spectra minus that of the untreated group to present the drug-mediated difference in spectral intensity. For 5Fu-treated cells, the subtraction results were mostly negative values for both 24h and 48h(figure6.a, b). The main decrease occurred at peaks of 1658 cm^-1^(Amide I, proteins) 1331 cm^-1^(collagen),794 cm^-1^(DNA),1092 cm^-1^(DNA), 1247cm^-1^(Amide III, protein) 1568 cm^-1^(Amide I, protein) 1334 cm^-1^(protein) due to 5Fu inhibition of thymidylate synthase and incorporation of its metabolites into RNA and DNA and mainly targeting at S phage^27^. As for the CKI-treated (2mg/mL) cells, the subtraction results of cytoplasm spectra were different from each other, which mainly expressed a negative value at 24h and a positive one at 48h. For 24h, almost all component was decreased which was reflected by the lower intensity at 1660 cm^-1^(acyl chain), 1576 cm^-1^(DNA), 1448 cm^-1^(Lipids/proteins), 1332 cm^-1^(Proteins), 1304 cm^-1^(Lipid/protein)1252 cm^-1^(Cytosine/adenine), 1122 cm^-1^ (proteins/lipids) and 1003cm^-1^(Phe vs(CC)ring). Interestingly, the peak intensity at 843 cm^-1^ was increased compared to phospholipids, which suggested that the increased cell activities related to the membrane. It was consistent with the strong uprising lipids signals at 48h in peaks 1743 cm^-1^(COOR),1436 cm^-1^ (acyl chain), 1077 cm^-1^ (Typical phospholipids). We infer that CKI might facilitate cell activities or pathways involving lipids/ fatty acids. The increasing of lipids signal and the disappearance of RS signals near 1745 cm^-1^ indicated the increase of lipids signal mainly resulted from the increase of fatty acid in the cytoplasm. Lipids are required to maintain cellular structure, supply energy, and involved in cell signaling. Lipid metabolism participates in the regulation of many cellular processes such as cell growth, proliferation, survival, apoptosis, autophagy.^42, 43^ Thus, related lipids/fat acid alterations could be one of the CKI anticancer mechanisms, considering the key role autophagy played in lipid metabolism and balance in cells and the cytotoxicity of free fatty acid.^44-46^

## CONCLUSIONS

Through a sub-cellular Raman spectroscopic analysis at the resolution of IVs, we captured multi-level cell changes caused by multi-targeted drug CKI. CKI caused DNA replication/repair inhibition was reflected by nucleic acid-related peaks decrease at cell nucleus RS. The enormous intracellular vesicle accumulation related to CKI induced apoptosis and autophagy was observed. The RS from intracellular vesicles and cytoplasm displayed the cell component degradation process meditated by IVs. The lipids/fat acid alterations showed that the lipids metabolism role in CKI anticancer effect and the mechanism needs to be further investigated. In general, we proved sub-cellular Raman spectroscopy is a powerful tool to explore the internal cell complexity, especially for a multi-target drug investigation.

## ASSOCIATED CONTENT

## Supporting Information

Assignments of Raman bands in spectra for cells (Table S1). Nucleus Raman spectroscopy at 48h (Figure S1). Cytoplasm Raman spectroscopy at 72h (Figure S2) (PDF)

## AUTHOR INFORMATION

## Author Contributions

All authors have given approval to the final version of the manuscript.

## Notes

The authors declare no competing financial interest.

## ACKNOWLEDGMENT

This work was supported by National Natural Science Foundation of China (NSFC) (U19A2007,32150026).

---

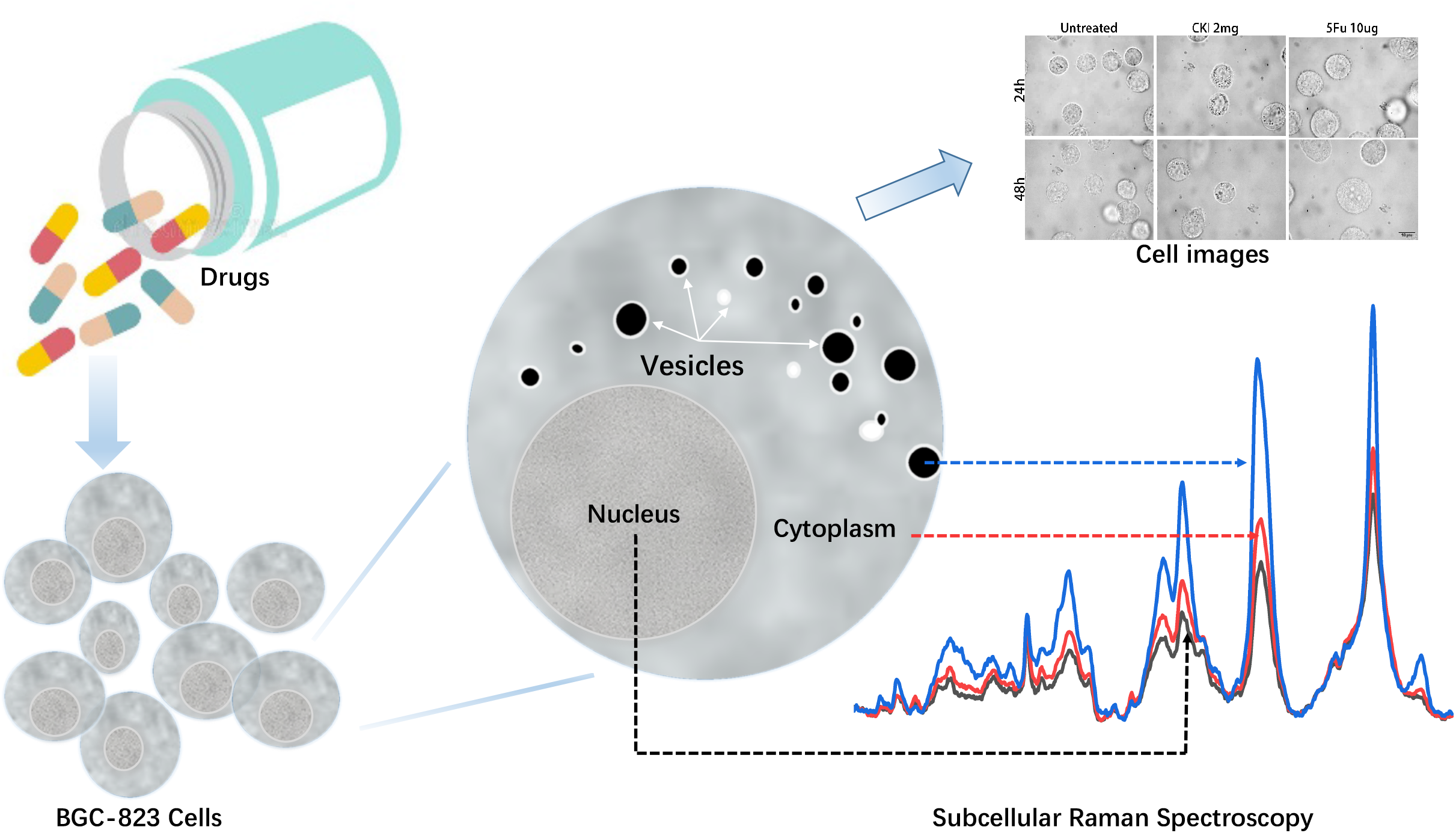

