## Supplementary material for "Sub-cellular dynamic investigation of the multi-component drug on the gastric cancer cell BGC823 using Raman spectroscopy"

### Table S1 Raman peak Assignment[1-17]

| Raman band(cm-1) | Assignment | | | |
|---|---|---|---|---|
| | Nucleic acids | Proteins | Lipids | Carbohydrates |
| **792** | DNA (ν(OPO)backbone) | | | |
| **780-805** | C,T,U (ν(CC)ring) | | | |
| **843** | Phosphodiester νs(O–P–O) | | | |
| **941** | RNA/ribose (ν(CC)ring) | | | |
| **1003** | | Phe (νs(CC)ring) | | |
| **1030** | | Phe | ν(CC),phospholipids | (δ(CH))ν(CC), ν(CO), ν(C–OH) |
| **1077** | ν(PO2−) | | | |
| **1092** | DNAν(PO2−) | | | |
| **1122** | | ν(C-N) ν(C-C) | ν(C-N) ν(C-C) | |
| **1199** | ν(PO2−) | ν(C-N) | | |
| **1252** | | ν(C-N) | | |
| **1296** | | amide III (δ(NH), ν(CN)) | | |
| **1313** | | | t(CH3CH2) | |
| **1334** | | ω(CH3/CH2) | | |
| **1375** | A,G,T (ν(CC)ring) | glycoproteins (δ(CH3)) | lipids/acyl chains (δ(CH3)) | saccharides |
| **1446** | | | lipid denaturation | |
| **1436** | | δ(CH2, CH3) | δ(CH2, CH3) in acyl chain | |
| **1579** | Guanine; adenine | | | |
| **1660** | | AmideI(ν(C=O))/α-helix | ν(C=C) | |
| **1720-1750** | | | ν(C=O)in ester COOR | |

A – adenine; G – guanine; Glu –glucose; Phe – phenylalanine; δ – in-plane deformation; γ – out-of-plane deformation; ν- stretching; ρ– rocking; t – twisting; u –wagging. s – symmetric

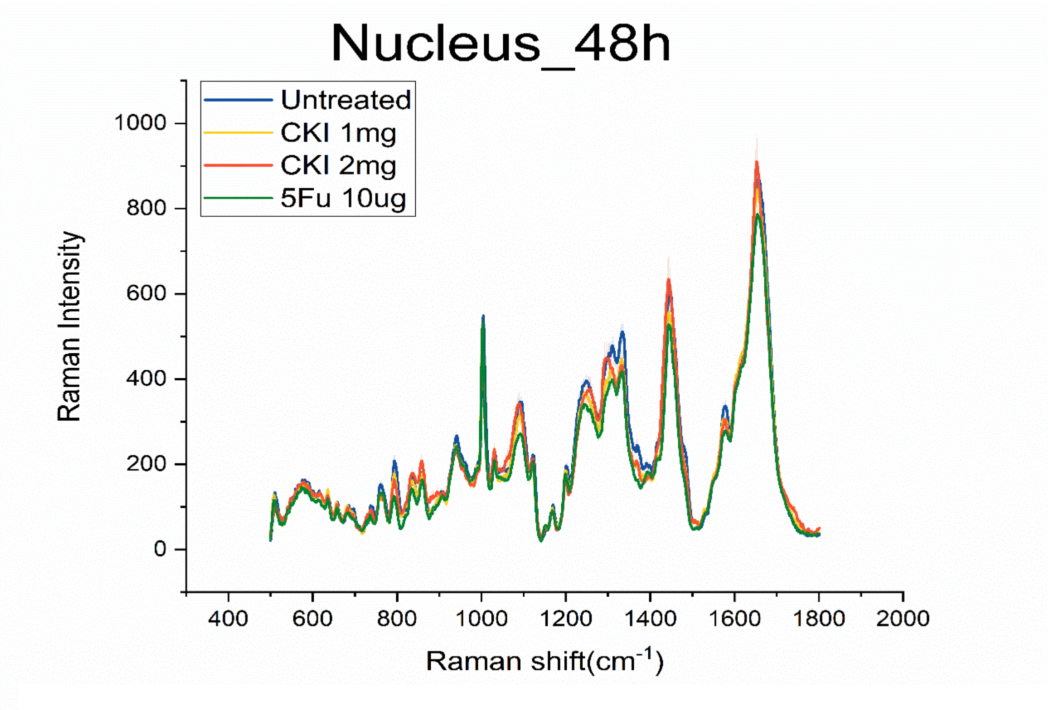

Figure S1. Nucleus Raman spectroscopy at 48h (mean ±s.e.m., the light shadow represents the s.e.m)

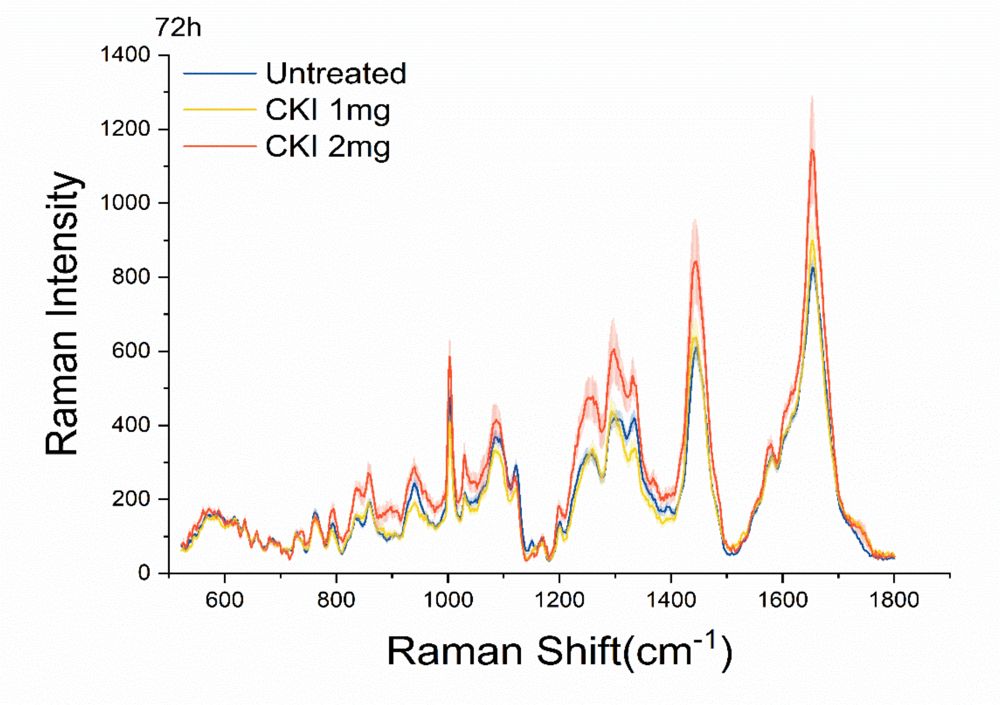

Figure S2. Cytoplasm Raman spectroscopy at 72h (mean ±s.e.m., the light shadow represents the s.e.m)

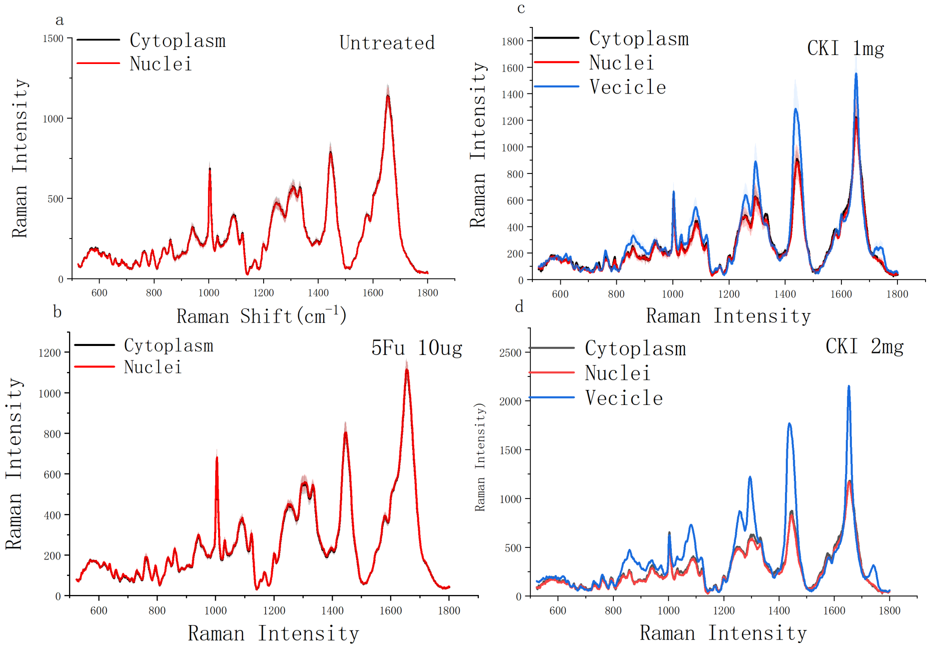

Figure S3. Subcellular RS of BGC-823 cells at 24h. a, d) The sub-cellular RS of the cells without drug-treatment; b) The image and sub-cellular RS of the cells under 5Fu 10ug/ml; c d) The sub-cellular RS of the cells under CKI 1mg/ml and CKI 2mg/ml treatment.

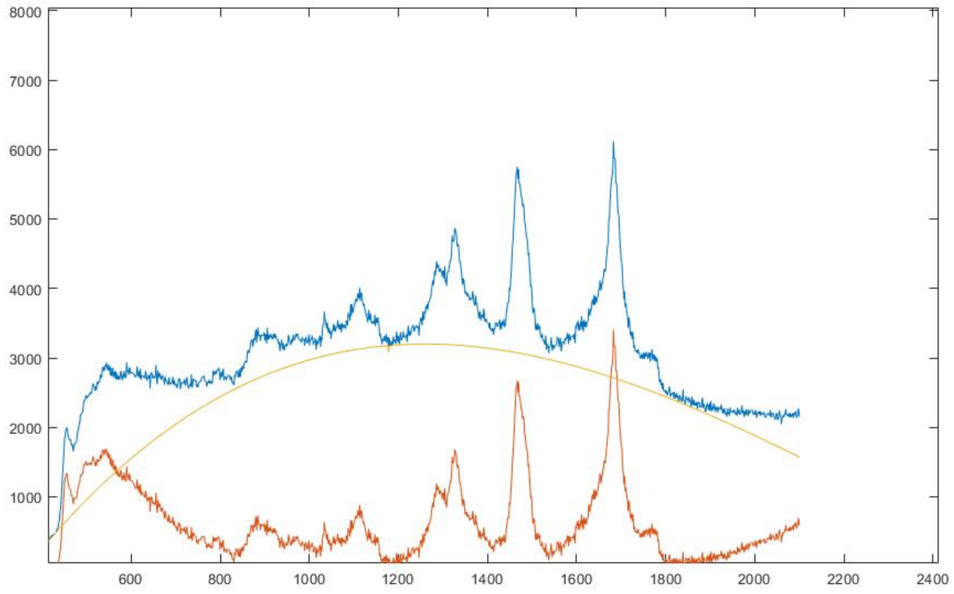

Figure S4. Spectrum before and after 3rd order polynomial .Blue line represents pre-process, Orange is after 3rd order polynomial. Yellow line is the 3rd order polynomial line

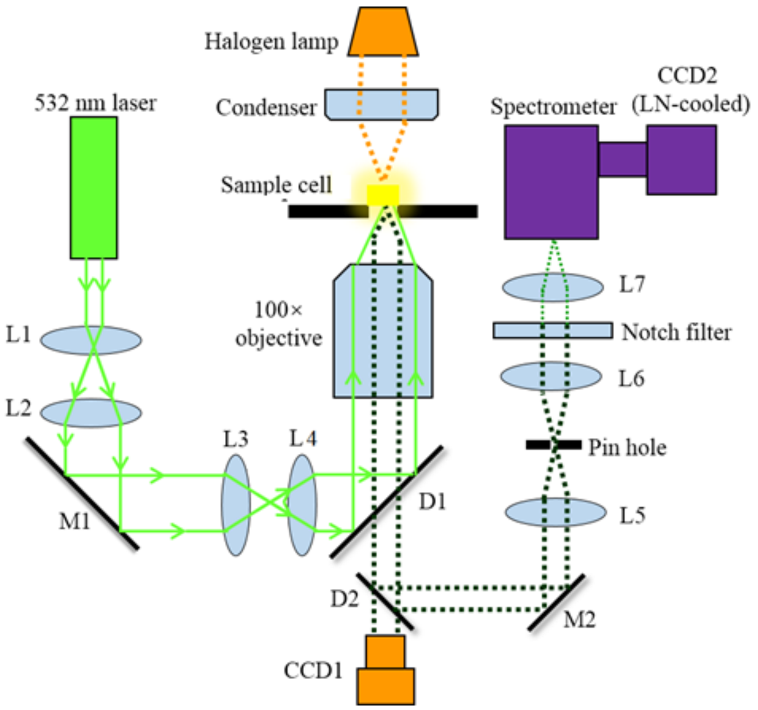

Figure S5. Configuration of the Raman spectroscopy
